# Neural responses during natural vision are action-timed rather than locked to the onset of stable foveal input

**DOI:** 10.1101/2024.10.25.620167

**Authors:** Carmen Amme, Philip Sulewski, Eelke Spaak, Martin N. Hebart, Peter König, Tim C. Kietzmann

## Abstract

Visual processing is traditionally studied using static viewing paradigms in which researchers analyse brain responses to the onsets of a sequence of randomly selected stimuli. Translating this “stimulus onset” approach to active vision paradigms that allow for free eye-movements, researchers often consider fixation onsets as the events that trigger visual activity across the visual system. Here, we test this assumption by analysing a large-scale magnetoencephalography (MEG) dataset with simultaneously recorded eye movements of 5 participants who freely explored thousands of natural images, yielding approximately 200,000 gaze events. We show that saccade-related events, particularly peak saccade curvature, rather than fixation onsets, explain most variance in latency of the early sensory component M100. Further, comparing the classic M100 elicited by stimulus onsets with the M100s elicited during active vision revealed stark differences, both in response to saccade-related and fixation-onset events. This indicates that phenomena discovered using gold standard stimulus onset paradigms do not necessarily translate to natural vision. Our findings challenge the prevailing approach to studying vision in static paradigms and highlight the importance of internally generated and action-driven signals in the dynamics of natural sensory processing.

## Introduction

To explore our environment, we move our eyes multiple times per second. Natural vision is therefore an inherently active process that samples input through a continuous sequence of saccades and fixations. In contrast, visual processing in the laboratory has long been studied with static experimental paradigms in which participants maintain central fixation while they are presented with discrete stimuli. In these classic settings, neural data are analysed relative to stimulus onset. As the field increasingly expands from static viewing toward more naturalistic active vision paradigms, this logic has largely carried over to free viewing. Since fixation onset marks the moment at which the eyes settle and visual input becomes relatively stable, it is often treated as the active-vision analogue of stimulus onset. Accordingly, fixation onset has commonly been used as the critical event for analysing neural responses during natural viewing ^[1–3]^.

However, active vision is not simply a series of passive visual snapshots. It is a self-generated and planned process in which each eye movement is embedded in an ongoing sequence of selection, preparation, and execution. During visual exploration, upcoming saccades are selected on the basis of peripheral visual information, through bottom-up saliency mechanisms ^[4]^ as well as top-down influences related to task demands and behavioral goals ^[5–8]^. Once a target has been selected and before saccade initiation, attention shifts from the current fixation location to the next ^[9,10]^. More generally, active vision relies on a continuously maintained representation of the environment that extends beyond what is currently foveated ^[11–14]^. Given this predictive and self-initiated structure of visual exploration, fixation onset need not be the defining temporal anchor of neural responses. Neural processing may instead depend on other eye-movement-related events and may begin before fixation onset itself ^[15]^. Determining which time point best captures the initiation of visual responses during natural viewing is, therefore, critical for understanding active vision.

Here, to test whether fixation onset is the appropriate event for aligning neural responses in active vision, we combined magnetoencephalography (MEG) and eye tracking during free exploration of natural scenes. Our results show that the M100, related to the well-studied P100 component in EEG, is initiated earlier than fixation onset and is more strongly related to saccade-related events, especially the time point of peak saccade curvature, a previously largely ignored active vision event.

We next asked a broader question, namely whether neural responses obtained in classical static viewing paradigms are comparable to those observed during active vision. To address this, we compared event-related fields (ERFs) locked to stimulus onset with ERFs locked to fixation- and saccade-related events during natural viewing. ERFs aligned to fixation- or saccade-related events showed distinct topographies from stimulus-onset ERFs, indicating qualitative differences in the underlying neural responses.

Together, these findings suggest that neural processing during active vision is not adequately captured by the event structure inherited from static paradigms. Instead of relating to purely external, stimulus-related signals, such neural processing is deeply intertwined with internally generated, eye-movement-related signals.

## Results

### M100 latency shifts with saccade duration

As a first approach to the question of which event better explains cortical responses during active vision, we compared ERFs locked to fixation-onset events and saccade-onset events in relation to various saccade durations. To be conservative towards fixation events, we selected, for each participant, the sensor with the highest amplitude in the fixation-locked response within 60 - 110 ms after event onset. We then computed the ERFs across 160 equally sized bins based on the preceding saccade duration (Fig. 1A). We found that the M100 deflections were better aligned with the preceding saccade onset than fixation onset (Fig. 1A). The same observation held when selecting sensors based on the saccade onset-locked ERF (Fig. 1B). To quantify this observation, we obtained ERF latency by determining the halfway point of the ERF leading to the peak of the M100. This observation was further supported by the finding of a negative relationship, a mean slope of −0.741 (standard deviation (SD) = 0.051), between the M100 latency and the saccade duration (Fig. 1C). A direct comparison between fixation and saccade onset ERFs for different saccade durations demonstrated much-improved alignment of saccade onset-locked ERFs (Fig. 1D). The finding that the main ERF deflections track saccade-related events in time, rather than fixation onsets, indicate that fixation onsets may not represent the underlying cause of the ERF.

**Figure 1.**
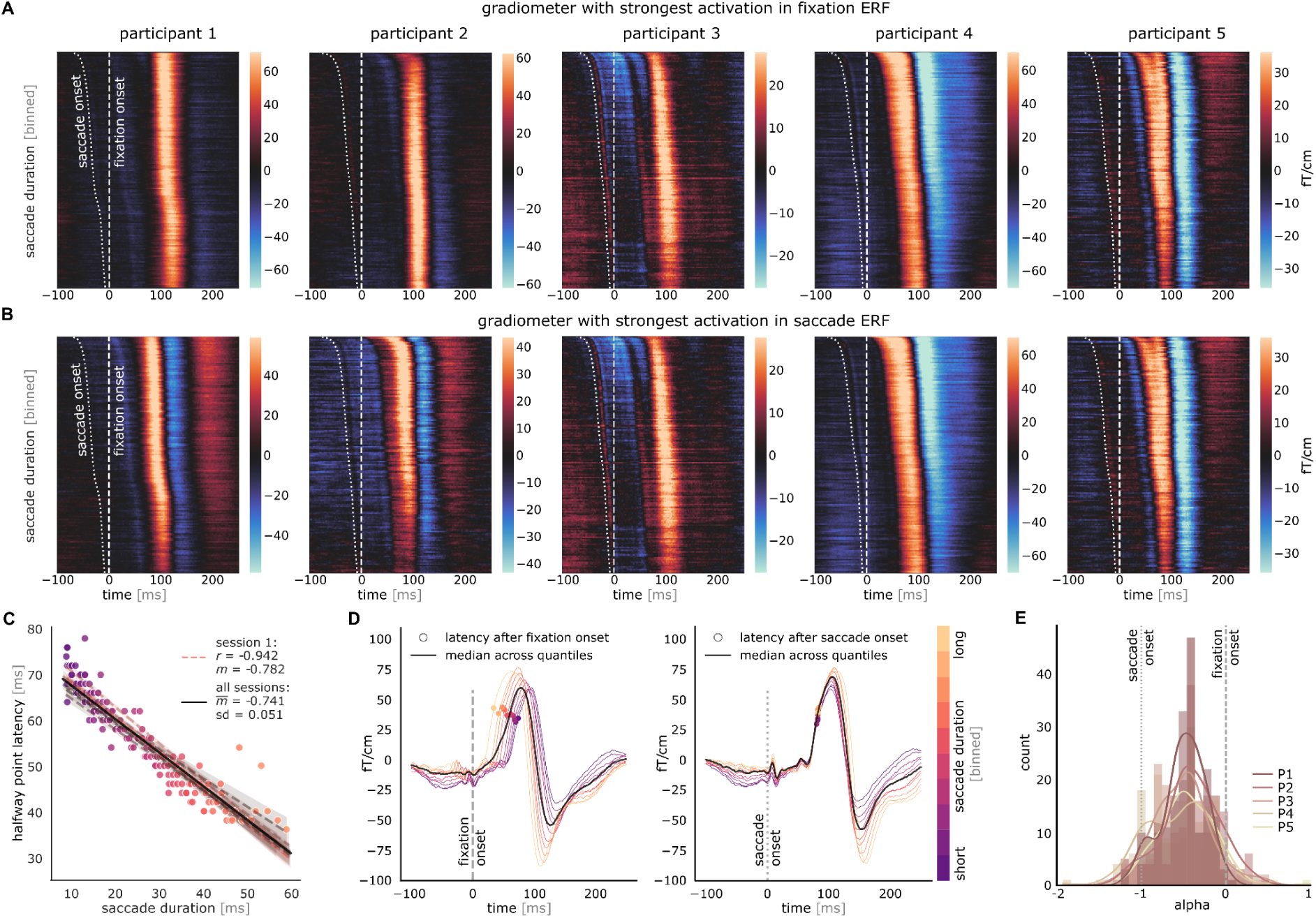
ERFs are locked in between fixation and saccade onset. **(A, B)** The saccade duration binned ERF deflections followed the onset of preceding saccades. **(C)** Session-wise linear fit for the previously selected sensor for participant 4 between the duration of the preceding saccade and the latency of the fixation onset-locked ERF per saccade duration bin. The ERF latency is defined as the halfway point of the slope leading to the M100 (see dots in D and E; with 95% CI). **(D)** The fixation (left) and saccade (right) onset-locked ERF per saccade duration bin (*n_bins_*= 10). **(E)** Across all participants and gradiometers that show a locking at any point in time, the most optimal ERF alignment for inter- and extrapolated event onsets around saccade (*α* = −1) and fixation onset (*α* = 0) is in between the onset of these two events.

### A parametric search for the optimal event latency shows a broad distribution in between and slightly beyond saccade and fixation onset

The slope between M100 latency and saccade duration fell between −1 and 0 (*m̅* = −0.741; Fig. 1C), suggesting that responses across the visual system during active vision may also be driven by events other than the previously explored fixation- and saccade-onset events. We therefore expanded our analyses to include hypothetical event onsets by shifting the reference event to before, in between, and after saccade and fixation onset. We parameterized this shift using *α*, where −1 is saccade onset and 0 is fixation onset (see *Methods*). For each saccade duration bin ERF, we marked the point of half-maximum on the ascending flank of the M100. For each gradiometer, we then identified the *α* value that best aligned the M100 half-rise points across saccade duration bins yielding the best ERF alignment score. We focused subsequent analyses only on gradiometers showing a well-defined alignment, meaning that the ERF alignment score exhibited a clear dip at some point in time as a function of *α*. We found that, for all participants and gradiometers included here, the distribution of optimal *α*-values across selected gradiometers indicates that optimal event latencies fall between saccade onset and fixation onset (Fig. 1E).

### The M100 is predominantly locked to peak saccade curvature

Because our alignment shifting analysis is agnostic to the identity of an event that could be triggering the ERF in the brain with the observed timing, we next compared several well-defined saccade- and fixation-related events. These include, in addition to saccade onset and fixation onset, peak saccade velocity, peak saccade curvature, peak visual motion energy, and post-saccadic oscillation offset. Fig. 2A shows the schematic time points of the saccade-related events on a real example saccade trajectory. To estimate visual motion energy, we constructed saccade movies from millisecond-resolved gaze positions across the respective scene images and processed using several spatiotemporal filters ^[16]^. Peak motion energy was then defined as the time point of the maximal total filter response (Fig. 2B). Relative to saccade onset, saccade velocity typically peaks first, followed by visual motion energy and saccade curvature. Visual inspection suggests that the timing of saccade-related events varies with saccade duration, whereas the offset of the post-saccadic oscillation appears unaffected (Fig. 2C).

**Figure 2.**
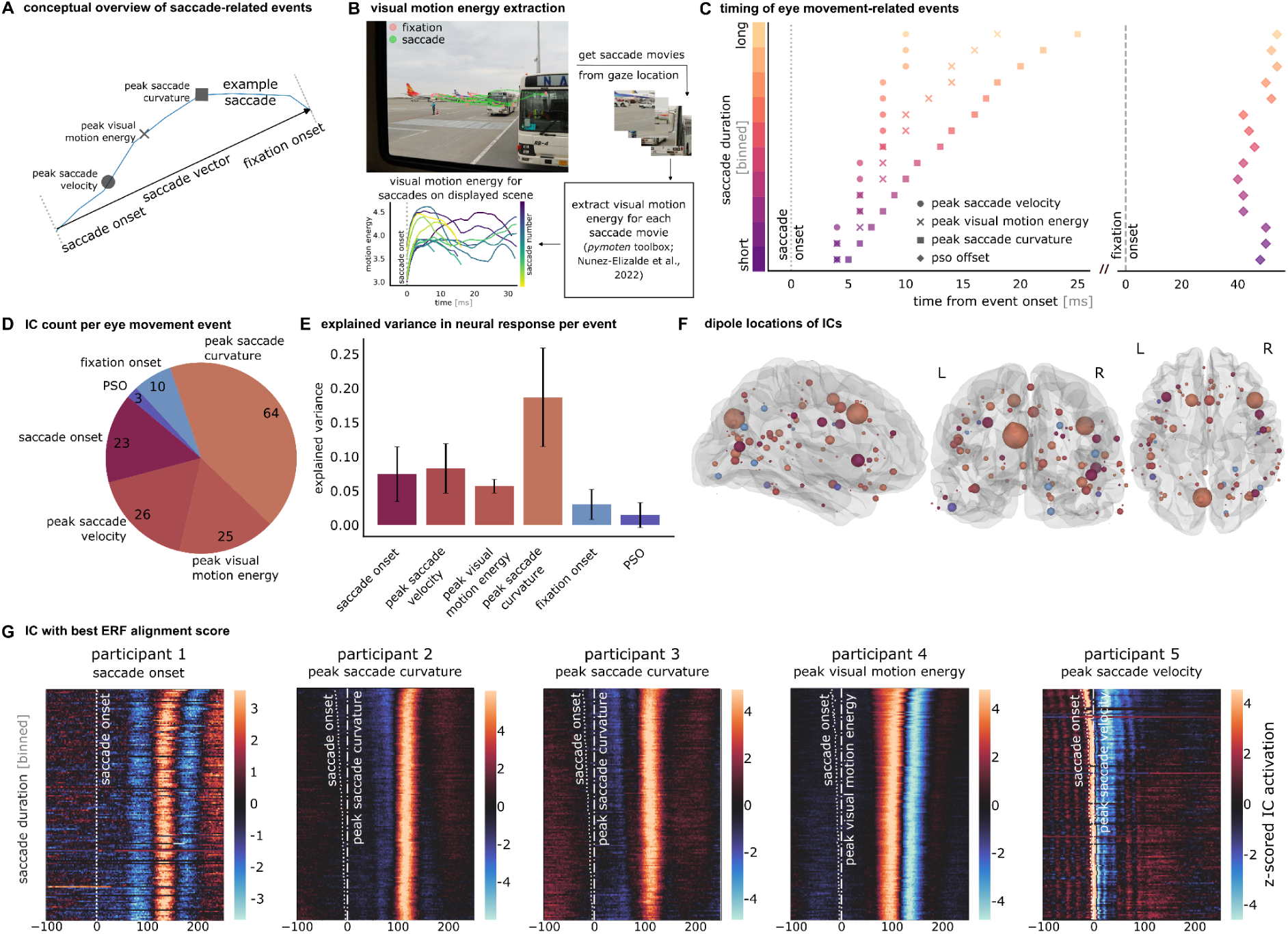
ERF alignment is best at the time point of peak saccade curvature. **(A)** Conceptual overview of a real example saccade trajectory with the schematic temporal onsets of saccade-related events. **(B)** For each saccade, we constructed a saccade movie by extracting image crops centered on the gaze location, covering 3° of visual angle, and sampled at 1000 Hz. Motion energy was subsequently computed using the *pymoten* toolbox ^[16]^, resulting in a temporal motion energy profile for each saccade. **(C)** Onset of eye movement-related events per saccade duration bin (*n_bins_* = 10), averaged across participants. **(D)** Per independent component (IC; n = 80 per participant), we determined the event with the best ERF alignment across saccade duration bins and accumulated the number of ICs across participants for each event. **(E)** Accumulated explained variance of the neural signal based on ICs associated with each event, averaged across participants, with 95% confidence interval around the standard error of the mean. **(F)** Dipole locations of ICs across participants, color-coded based on the event that aligns saccade duration-binned ERFs best, and scaled based on explained variance in the neural signal. **(G)** IC with the best ERF alignment score per participant, locked to the onset of the optimal event.

As a second change to the previous analysis approach, we decomposed the MEG signals into independent components (ICs; 80 ICs per participant), because the measured signals at the sensor level can mirror multiple overlaid brain processes with different event locking. Applied to each participant’s neural data, this allowed us to identify the relation of each component to the different gaze events described above. ICs for which the dipoles were clearly located outside of the brain were discarded from further analyses (across participants, 6.8 ICs were discarded on average). To determine ERF alignment, we divided the neural data of each session into 15 saccade-duration bins, locked the IC activity to all event types respectively, and obtained the ERF alignment score across saccade duration bins. An ANOVA was conducted across sessions to select ICs showing a significant modulation of peak latency by event type. For all significant ICs, we determined the optimal event by selecting the IC with the best (i.e., lowest) median ERF alignment score. Out of the 151 significant ICs across participants, only 8.6% of them showed optimal ERF alignment to fixation-related events, whereas 91.4% of the ICs showed optimal ERF alignment at saccade-related events, particularly at the point of peak saccade curvature (42.4%; Fig. 2D). ICs aligned to saccade-related events explained the largest proportion of neural signal variance (40.0%, 95% CI [310%, 49.1%.]), whereas fixation-related events contributed minimally (4.5%, 95% CI [1.6%, 7.3%]; mean across participants; Fig. 2E). At the event level, peak saccade curvature accounted for the largest share of IC-explained neural variance (18.7%, 95% CI [11.5%, 25.9%]). Finally, to assess the spatial distribution of these ICs across cortex, we examined their dipole locations. We found that across participants, the dipole locations of the ICs corresponding to their optimal event are spread across large parts of the cortex, suggesting that active vision on natural scenes recruits neural sources beyond visual areas (Fig. 2F).

### Stimulus onset differs from both peak saccade curvature and fixation onset

We next turned to a more general question about the ability to generalise from passive to active vision paradigms. In particular, we asked whether stimulus onset ERFs, the most common approach towards studying vision, exhibit similar topographies to ERFs computed in response to active vision events. We considered peak saccade curvature (the event type with best ERF alignment scores in the largest number of ICs in our previous analysis) and fixation onset events. A first visual inspection indicates clear differences in the resulting topography of the scene onset ERF and from fixation onset and peak saccade curvature ERFs (Fig. 3A). To account for potential differences in timing across the ERFs, we applied Dynamic Time Warping (DTW; Itakura, 1975; Müller, 2007; Sakoe & Chiba, 1978) separately to each participant and sensor to obtain a latency-corrected ERF dissimilarity score (Fig. 3B illustrates an example for one participant and one sensor). When averaging dissimilarity scores across magnetometers and participants, it becomes apparent that scene onset ERFs are different to ERFs computed by both fixation onsets and saccade curvature locking (Fig. 3C). Mean dissimilarity was highest for scene ERF versus peak saccade curvature ERF (*M* = 4.77, SD = 3.33) and scene ERF versus fixation ERF (*M* = 4.25, SD = 2.87), whereas fixation ERF versus peak saccade curvature ERF yielded substantially lower scores (*M* = 0.61, SD = 0885). FDR-corrected sensor-wise paired t-tests supported this pattern. Only 2 sensors differed significantly between scene ERF versus fixation ERF and scene ERF versus peak saccade curvature ERF, whereas fixation ERF versus peak saccade curvature ERF differed from both scene-based comparisons much more strongly, with 42 significant sensors relative to scene ERF versus fixation ERF and 65 relative to scene ERF versus peak saccade curvature ERF. The corresponding dissimilarity matrix further shows that any comparison involving the scene onset ERF produces higher scores, whereas the dissimilarity between fixation and peak saccade curvature is of comparable magnitude to the split-half within-event comparison along the diagonal (Fig. 3D). Consistent with this pattern, only a single sensor showed a significant difference between the scene ERF and peak saccade curvature ERF split-half within-event comparisons.

**Figure 3.**
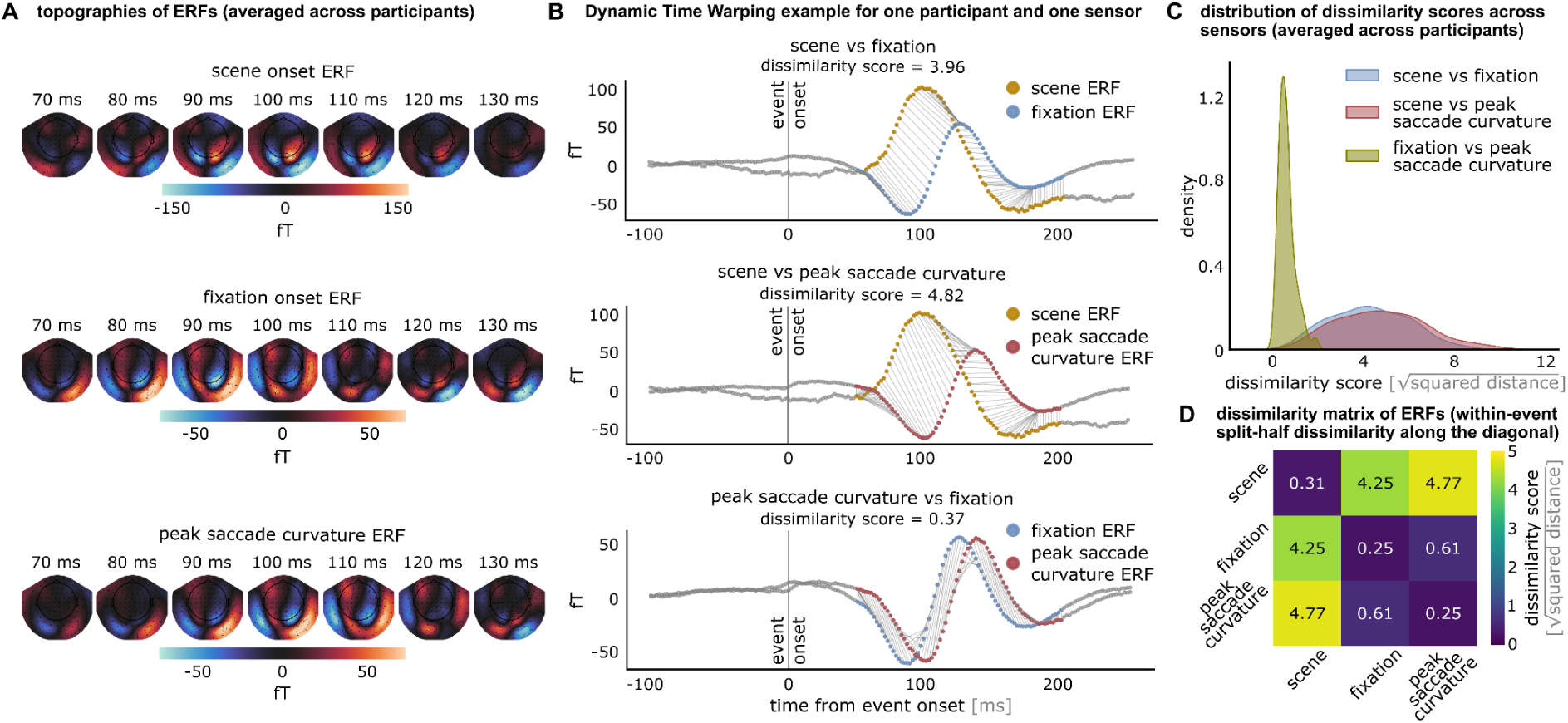
Spatiotemporal differences between stimulus onset and fixation/peak saccade curvature ERFs. **(A)** Topographies of scene, fixation and peak saccade curvature onset-locked ERFs, averaged across participants. **(B)** Example of Dynamic Time Warping (DTW) for one sensor and participant to compare the 3 different event onset-locked ERFs, applied between 50 and 200 ms after event onset. The dissimilarity score 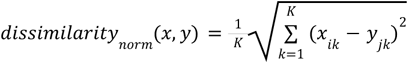, where *x* and *y* are the two time series and *K* the length of the warping path) is obtained for each point after alignment. **(C)** Distribution of sensor-wise DTW dissimilarity scores per event comparison, averaged across participants. **(D)** Dissimilarity matrix showing the average DTW dissimilarity across participants and sensors. Trials of the same event type along the diagonal were split in half, and the number of observations was matched across splits to compute within-event dissimilarity.

## Discussion

By analysing data from a large-scale MEG and eye tracking dataset across thousands of natural images, we here tested a central assumption in active vision research: that fixation onsets are the most appropriate reference event for explaining MEG responses recorded during natural viewing. Contrary to this assumption, we find that fixation onset is not the main event to which the MEG responses are best aligned. Although earlier EEG, iEEG, and macaque electrophysiology studies showed that fixation-locked neural responses are modulated by saccade duration ^[20–22]^, none of these studies systematically investigated whether other saccade-related events best account for this modulation. Here, using a large-scale MEG dataset during natural vision in humans, we find that saccade-related events, predominantly the time point of peak saccade curvature occurring shortly after saccade onset, provide the strongest alignment of the neural responses. These findings highlight the importance of internally generated processes during natural vision that are initiated well before fixation onset, and support a view of sensory processing as fundamentally action-driven.

Importantly, our results furthermore demonstrate that neural responses to stimulus onsets, the gold standard in vision research, differ significantly from MEG data in response to naturalistic gaze events (both peak saccade curvature and fixation onset ERFs). This finding is consistent with a previous study showing that the sink-source configuration in non-human primate V1 is oriented in the opposite direction between stimulus and fixation onset ^[23]^. This observation suggests qualitative differences between stimulus-onset driven analyses and natural vision. The key distinction between stimulus-onset events and events related to eye movements is that eye movements are planned and self-initiated, whereas the temporally and semantically unexpected onset of a stimulus is not. Self-generated eye movements have been shown to influence neural responses in early visual areas ^[24]^ and may stem from internally generated signals related to predictions of visual information within the predictive coding framework ^[25–27]^, contextual ‘priming’ ^[28,29]^ and memory ^[30]^, or they may stem from a ‘readiness potential’ signalling the visual system to prepare for the incoming information ^[31,32]^. Future work is needed to investigate these or other possibilities.

Notably, we observed that the largest number of ICs exhibited event-related responses that were temporally locked to peak saccade curvature, an event not commonly considered in terms of MEG response locking during active vision. Experimental work has linked saccade curvature to competitive interactions between target and distractor representations in oculomotor structures, where the direction and magnitude of deviation depend on factors such as saccadic reaction time, prior knowledge, distractor salience, and spatial layout ^[33–41]^. These competitive dynamics have been proposed to arise from time-varying changes in the relative activation of target and distractor representations across the superior colliculus and frontal eye fields, though the precise contributions of top-down inhibition, lateral interactions within oculomotor maps, and brainstem feedback remain debated ^[34,35,42–44]^. The temporal locking we observe may thus reflect the resolution of such competitive processes, suggesting an interplay of motor planning, attentional modulation, and stimulus-driven activation that shapes the latency of eye movement-related neural activity. This new perspective on the M100 during active vision opens a new avenue of research only accessible when studying vision in its natural, active form.

Taken together, our results imply that neural processing in natural vision is guided by an internally generated signal, whereby self-initiated actions are a key to sensory processing. Studying active visual paradigms, both in terms of experimental data and computational modeling, hence seems unavoidable for understanding visual information processing as it occurs naturally in everyday settings.

## Methods

### The MEG - Eye-Tracking Data Set

#### Procedure

Each of the 5 participants completed 10 measurement sessions over a period of 2 - 5 weeks. They had to freely visually explore images and to give a verbal caption in 25% of the trials, which they were prompted to do after image presentation. The caption and non-caption images were presented in random order. The participants were seated with a viewing distance of 70 cm to the screen, and the stimuli were presented with a visual angle of 28.54° x 21.61°. At the beginning of the first session, all participants were given instructions in German. The participants were then shown 10 example images with 5 example captions per example image. At the beginning of each session, the eye tracker was calibrated using a 9-point calibration where participants were asked to fixate sequentially appearing dots at 9 different locations on the screen. Every new trial began with the central appearance of a white fixation cross on a gray background for a minimum duration of 1s, which also served as a drift control for the fine-calibration of the eye-tracking system. If the fixation deviated more than 1.3° from the fixation cross, the trial was restarted. All stimulus images were displayed for 4s, allowing participants to freely explore them through natural eye movements. On 75% of the trials, the presentation of the fixation cross indicated the start of the subsequent trial. In the remaining 25% of the trials, the stimulus presentation was followed by the appearance of a black microphone on a gray background for 1s in the center of the screen, prompting the participants to give a verbal description of the images. The recording of the caption started with the disappearance of the microphone symbol and ended after a duration of 8s with the start of the following trial. The stimuli were presented in blocks of 30 trials. The first session consisted of 10 blocks with 300 trials in total. The other 9 sessions consisted of 14 blocks each (420 trials/session). After every 3 to 4 blocks, the eye tracker was recalibrated, and the participants could take a small break. After the experimental sessions, the recordings of the captions were transcribed manually.

#### Eye Tracking Data Acquisition and Preprocessing

Eye tracking data were recorded using a SR Research EyeLink 1000 system (SR Research; https://www.sr-research.com) with a sampling rate of 1,000 Hz. The raw eye-tracking data were preprocessed through an algorithm by Engbert & Mergenthaler (2006) to detect saccades. Velocity vectors were obtained by taking the sample-wise differences, separately for x- and y-coordinates. The x- and y-velocity vectors were scaled by dividing by the standard deviation. Speed was then calculated by taking the norm of the sample-wise velocity vectors. Samples that exceeded 5 standard deviations were classified as saccades. Blinks were automatically detected by the eye tracker. All samples that were not classified as a blink or a saccade were classified as a fixation. Because we were only interested in fixations that finished during the stimulus presentation, the last fixation on all images was excluded.

#### MEG Data Acquisition and Preprocessing

To minimize head movements during the measurement, we created individually fitted head stabilizers for each participant. We took each participant’s head shape using a Structure Sensor Pro (XRPro, LLC; https://structure.io) and compared that to a 3D scan of the MEG head piece. The space between the MEG headpiece and the participant’s head shape was milled into styrofoam. Before each measurement session, a Polhemus FASTRAK tool (Polhemus; https://polhemus.com) was used to digitize the participants’ head shape. To track head positions inside the MEG, a total of five coils were placed above the nasion on the forehead, at the inions bilaterally, and in between these points on the forehead. The experiment took place in a magnetically shielded room using a NeuroMag VectorView MEG system (Megin; https://megin.com) consisting of 306 channels (102 magnetometers, 204 planar gradiometers). The raw data were recorded with a sampling rate of 1,000 Hz, and an online filter of 0.1-330 Hz was applied.

After the recording, the raw MEG data was bandpass filtered through a causal bandpass filter between 0.2 and 200 Hz and downsampled to 500 Hz. MEG channels that were visually observed to be noisy during the measurement were tracked and interpolated before analysis. The raw MEG data was then cleaned by applying a spatiotemporal *MaxFilter* (maxwell filter; Elektra), including a temporal Signal Space Separation (tSSS; Taulu et al., 2004), which also accounted for different head positions and head movements made during the measurement.

All further analyses were implemented in Python by making use of the toolbox MNE-Python ^[47]^ and several custom written scripts. We applied independent component analysis (ICA) separately for each participant and session to identify and remove noisy components. To identify and remove eye movement-related components, we correlated the epoched fixation-based data of each component with the corresponding x-/y-coordinates of the eye position. We median-centered the data per channel and session to account for different session-wise offsets in the MEG data.

#### MEG-ET Alignment

To investigate fixation-related neural processes, we mapped the MEG data to the eye tracking data by aligning the MEG and eye tracking data to the scene onset, which was annotated in both the MEG and eye tracking data, and then mapped fixation annotations onto the MEG data based on fixation/saccade onset latency and duration within each scene presentation. The MEG data was then epoched based on fixation or saccade onset.

#### Participants

We recorded MEG data from 5 right-handed participants (3 female), all aged between 26 and 33 years (*m̅*_age_ = 27.8). All participants were native German speakers, reported to be healthy, and had normal or corrected-to-normal vision. Everyone gave written informed consent at the beginning of each experimental session, and additionally on the days for scanning and fitting the personalized head stabilizers. The study was approved by the ethics committee of the University Medical Center Leipzig and adhered to the Declaration of Helsinki. All participants received monetary reimbursement for their participation.

#### Stimuli

We included 4,080 images from the Natural Scenes Dataset (NSD; Allen et al., 2022), which consists of 73,000 images of natural scenes from the Microsoft Common Objects in Context (MS COCO) image dataset ^[49]^. We pseudo-randomly selected images from NSD with the following condition: To make sure that the semantic content of the images is equally represented, we performed k-means clustering on all NSD scene captions and identified 60 different semantic clusters. We selected an equal number of images from each cluster. Out of the 4,080 sampled images, 25% of the images (*n* = 1,020) were caption-images, where participants were asked to provide a verbal description, which were selected based on the following conditions: Allen et al. (2022) defined a set of 100 images that maximally span the semantic space (‘special100’), which were included as caption images. The ‘shared1000‘, which are images presented to all participants that were part of NSD, were also included as caption images, but only if the equal distribution of clusters allowed it. All participants were presented with the same caption and non-caption images, which were randomly shuffled for each participant.

### Data Analysis

#### Shifted Latency Analysis

To remove noisy epochs, we rejected epochs whose maximum amplitude exceeded the 99th percentile across all epochs. We additionally rejected epochs whose correlation with the median-based ERF fell below the 1st percentile across all epochs. This step was applied for all following analyses. The fixation-based epochs were binned into equally sized bins (n_bins_ = 160) based on the duration of the preceding saccade. For each saccade duration bin, we obtained a mean-based ERF. These ERFs were visualized in a heatmap showing the response of a single sensor sorted by the duration of the preceding saccade, with each row corresponding to one saccade duration bin (Fig. 1A, B). The participant-wise sensor selection was based on the strongest positive gradiometer activation between a time window from 60 to 110 ms of either the fixation-locked ERF (Fig. 1A; P1: MEG1923, P2: MEG1733, P3: MEG1933, P4: MEG2123, P5: MEG2512) or the saccade-locked ERF (Fig. 1B; P1: MEG2343, P2: MEG2512, P3: MEG1933, P4: MEG2123, P5: MEG2512). All further analyses described in this section were based on the sensor with maximum activation for saccade-locked ERFs.

For reliable halfway point detection, we inverted the polarity of the data whenever the peak ERF amplitude between 80 and 110 ms was negative. We defined ERF latency as the time point at which the signal reached the halfway point of the rising slope toward the maximum deflection. To reduce noise, ERFs were low-pass filtered using a central boxcar filter with a 30 Hz cutoff. The ERFs were cropped to a window spanning 80 ms before to 80 ms after the peak of the main deflection. For each saccade duration bin, the ERF halfway point was constrained to fall within a temporal window centered on the main ERF halfway point and extending ± the overall median saccade duration of that bin. Within this restricted window, the halfway point was defined as the midpoint between the peak amplitude and the minimum value of the main ERF within the 80 ms preceding the peak.

To investigate the relationship between saccade duration and ERF latency, we fitted a linear regression between saccade duration and the latency of the halfway point of the fixation onset-locked ERF. Latencies were binned based on the duration of the preceding saccade (n_bins_ = 160), and the regression was computed separately for each of the 10 measurement sessions (dashed lines; Fig. 1C). We obtained the correlation coefficient (r) and the slope (m) of the fitted line (reported only for session 1), as well as an average across all sessions.

To illustrate the shift of the ERF as a function of the preceding saccade duration, we binned the fixation and saccade ERFs into 10 equally sized bins defined by the duration of the previous saccade and visualized the mean-based ERF, either fixation onset-locked (Fig. 1D, left) or saccade onset-locked (Fig. 1D, right).

#### Mixing Factor Analysis

For each gradiometer, we binned the fixation epochs into 160 equally-sized bins based on preceding saccade duration and linearly inter- and extrapolated event onsets for each saccade duration bin. The simulated event onset can be defined as:

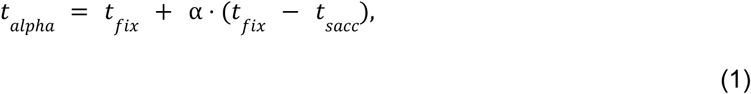

where *t_alpha_* represents the simulated event onset, α the temporal distance with respect to fixation onset, and *t_fix_* and *t_sacc_* the fixation and saccade onset time, respectively. A mixing factor of α = 0 corresponds to fixation onset (*t_fix_*) of a given epoch, whereas a mixing factor of α =− 1 indicates saccade onset (*t_sacc_*). A mixing factor of − 1 < α < 0 corresponds to an interpolated event onset between saccade and fixation onset. Accordingly, − 2 > α >− 1 indicates an extrapolated event onset up to twice the duration of the saccade before *t_fix_* ≙ α = 0, and 0 > α > 1 indicates an event onset up to the saccade duration after *t_fix_* ≙ α = 0.

As a measure of how well any α-based ERF could explain the single epochs, we obtained the standard deviation (SD) of the halfway point leading to the peak of the α-based ERF, separately for each gradiometer. The halfway point was determined as described in the paragraph above under *Shifted Latency Analysis*.

To only include gradiometers that show a clear preference for any event, we fitted a polynomial with a degree of 2 on the obtained alpha values, defined as:

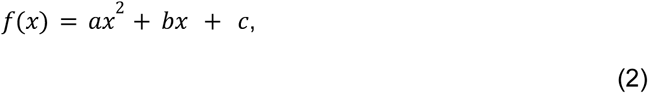

where *a* determines the width of the parabola. We performed a permutation test where we shuffled across α values (n shuffles = 1,000) to determine whether the *a*-value in the polynomial of the ordered α values is significantly higher (significance threshold = 0.05), indicating a clear dip in the parabola and a preference for an event onset. Fig. 1E shows the distribution of the α value with the lowest SD (i.e., best ERF alignment across saccade duration bins) of all gradiometers that are significant after fitting the polynomial, separately for each participant.

### Comparison of Different Eye Movement-Related Event Onsets

#### ICA and Dipole Fitting

To better isolate sources of neural activity, ICA was applied to all magnetometers of the scene onset-locked epoched neural data. Per participant, we obtained 80 independent components (ICs) and applied them onto fixation onset-locked neural data. To discard ICs that are related to noise, we applied dipole fitting on the ICs and excluded ICs where the corresponding dipole was fitted to a location outside the skull. On average, we discarded 6.8 ICs across participants.

#### Saccade Velocity

To obtain peak saccade velocity, we calculated the Euclidean distance between the current and the following eye tracking sample. To obtain the relative velocity with respect to the saccade vector, we computed the projection of the instantaneous velocity vector onto the saccade vector using the dot product, yielding the velocity component along the saccade direction:

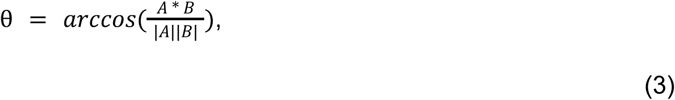

where *A* and *B* correspond to the sample and overall saccade vector, respectively. Relative saccade velocities were calculated at each ms.

For each session separately, the MEG data of all fixations was binned into 15 equally-sized bins according to preceding saccade duration, and the bin-wise data was locked to the timepoint of peak saccade velocity. To quantify the session-wise latency shifts of the 15 peak saccade velocity-locked ERFs, we obtained the SD of the halfway point leading to the peak of the M100 as described above under *Shifted Latency Analysis*, separately for each IC.

#### Post-Saccadic Oscillations (PSO)

Velocities were calculated at a sampling rate of 1,000 Hz from fixation onset on as described above (see *Saccade Velocity*). To obtain the average level of movement during a fixation as a baseline, we averaged velocities between 50 and 100 ms after fixation onset, which is later than the average PSO duration of 24 ms and before onset of the following saccade ^[50]^. PSO offset was defined as the first timepoint after fixation onset when the velocity falls within a window of +/− 1 standard deviation of the baseline:

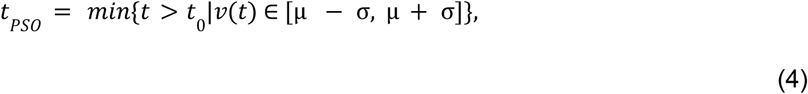

where *t*_0_ is fixation onset, *v* is velocity, µ is average velocity between 50 and 100 ms after fixation onset, and σ is the standard deviation of velocity between 50 and 100 ms after fixation onset.

For each session separately, the MEG data of all fixations was binned into 15 equally-sized bins according to preceding saccade duration. We then obtained the timepoint of PSO offset for each of the 15 saccade duration bins per session based on the median velocity per bin. To quantify the session-wise latency shifts of the 15 PSO-locked ERFs, we obtained the SD of the halfway point leading to the peak of the M100 as described above under *Shifted Latency Analysis*, separately for each IC.

#### Visual Motion Energy

At a sampling rate of 1,000 Hz, we extracted 100 × 100–pixel image patches corresponding to 3°, centered on the current eye position. This process yielded movies for each individual saccade. To obtain visual motion energy for each movie, we made use of the *pymoten* toolbox implemented in Python ^[16]^. We constructed spatiotemporal filters covering the range of spatial and temporal frequencies that the human brain is sensitive to. We constructed spatial filters between a range of 0.25 and 7 cycles/°, because early visual areas such as V1 respond to spatial frequencies up to almost 7 cycles/° ^[51,52]^ and MT to frequencies as low as 0.25 cycles/° ^[53]^. We then included 6 logarithmically evenly spaced spatial frequencies at 0.25, 0.5, 1, 2, 4 and 8 cycles/°. Because our longest saccade durations are about 60-70 ms, corresponding to 15 Hz, and MT responds to temporal frequencies up to 32 Hz ^[53]^, we decided to include 4 evenly spaced temporal frequencies at 15, 23, 32, and 40 Hz. Additionally, we also included 8 evenly spaced spatial filter directions at 0, 45, 90, 135, 180, 225, 270, and 315°. This resulted in a total of 4,800 spatiotemporal filters, tiled across spatial positions. The filters were then applied along the spatial and temporal dimensions of the saccade movie. The quadrature pair output of each filter was log-transformed before we averaged the outputs of the spatial filters to obtain a temporally resolved motion energy for each saccade movie (see Fig. 2B for a conceptual overview).

For each session separately, the MEG data of all fixations was binned into 15 equally-sized bins according to preceding saccade duration, and the bin-wise data was locked to the timepoint of peak visual motion energy. To quantify the session-wise latency shifts of the 15 peak visual motion energy-locked ERFs, we obtained the standard deviation of the halfway point leading to the peak of the M100 as described above under *Shifted Latency Analysis*, separately for each IC.

#### Saccade Curvature

For each saccade, we defined a straight saccade vector between saccade onset and offset and computed the perpendicular distance of each eye tracking sample during the saccade from this line ^[33,54]^. Saccade curvature was quantified as the maximum of these perpendicular deviations, normalized by saccade length to account for amplitude differences. The time at which this maximum deviation occurred was defined as the tim point of peak saccade curvature.

For each session separately, the MEG data of all fixations was then binned into 15 equally-sized bins according to preceding saccade duration, and the bin-wise data was locked to the timepoint of peak saccade curvature. To quantify the session-wise latency shifts of the 15 peak visual saccade curvature-locked ERFs, we obtained the standard deviation of the halfway point leading to the peak of the M100 as described above under *Shifted Latency Analysis*, separately for each IC.

#### Event Comparisons

To select ICs that show a preference for any event, we applied a one-way ANOVA across the SDs with the different events (saccade onset, peak saccade velocity, peak visual motion energy, peak saccade curvature, fixation onset, PSO offset) as groups and the 10 measurement sessions as observations. For each significant IC we assigned a best event based on the lowest median SD. In Fig. 2D we show the number of significant ICs per event. For Fig. 2E, we added the neural variance that is explained by each significant IC and averaged across participants, and plotted the explained neural variance with a 95% confidence interval line.

For Fig. 2.F, we transformed the dipole locations into *fsaverage* ^[55]^ coordinates and plotted all dipole locations across participants in a glass brain, color-coded based on the event that aligns saccade duration-binned ERFs best, and scaled based on explained variance in the neural signal.

For Fig. 2G, we selected the IC with the best ERF alignment score (i.e., lowest SD, obtained as described under *Mixing Factor Analysis*). We locked the data to the onset of the preferred event and binned the epochs into equally sized bins (n_bins_ = 160) based on the duration of the saccade. For each saccade duration bin, we obtained a mean-based ERF. The ERFs were then visualised in a heatmap.

### Scene Onset Analyses

#### ERF Topographies

To make sure we didn’t include eye movement-related data to the scene onset ERFs, the epochs were masked from the time point of the first saccade onset on. We then locked the neural data to either scene, fixation or peak saccade curvature onset and plot the ERF topographies, averaged across all participants, for a number of time points (Fig. 3A).

#### Dynamic Time Warping

Based on the ICA decomposition applied to the magnetometer data, as described above under *ICA and Dipole Fitting,* we removed ICs whose dipole locations were outside the skull, thereby excluding non-neural sources from the neural responses. To account for potential temporal shifts between ERFs, we applied Dynamic Time Warping (DTW) using the *DTAIDistance* toolbox in Python ^[56]^. To constrain the maximum temporal shift allowed between the two ERs, we set the DTW window size to 15 samples, corresponding to 30 ms given the 500 Hz sampling rate of the neural data. The DTW dissimilarity corresponds to the square root of the accumulated squared distances along the alignment path. To obtain a dissimilarity score after alignment, we divided this distance by the path length, thereby normalizing for the number of temporal shifts required to achieve optimal alignment. We performed pairwise DTW comparisons of ERFs elicited by scene onset, fixation onset, and peak saccade curvature, separately for each participant and magnetometer. For Fig. 3C, we averaged the resulting dissimilarity scores across participants for each magnetometer and plotted the resulting distribution. For the dissimilarity matrix shown in Fig. 3D, we averaged the pairwise dissimilarity scores across all magnetometers and participants. For the within-event comparisons along the diagonal of the matrix, we matched the number of trials per event, randomly split the trials into two halves, and compared the resulting ERFs within each event.

For the statistical analysis, we z-scored similarity scores within each subject across all sensors and comparisons. We then ran sensor-wise paired t-tests across subjects for each pair of event comparisons to test whether mean similarity scores differed between conditions at each magnetometer. The resulting p-values were corrected for multiple comparisons across sensors within each pairwise comparison using the Benjamini-Hochberg FDR procedure.

## Acknowledgements

This work was supported by the Deutsche Forschungsgemeinschaft (DFG, German Research Foundation) - 456666331. C.A., P.S., and T.C.K. are supported by the ERC Starting Grant TIME (101039524). P.S. is supported by the Federal Ministry of Education and Research (BMBF) and the Max Planck Society (MPG; Max Planck School of Cognition), and Deutsche Forschungsgemeinschaft (DFG; German Research Foundation) - GRK 2340. M.N.H. is supported by a research group grant by the Max Planck Society, the ERC Starting Grant COREDIM (101039712), the Hessian Ministry of Higher Education, Science, Research and Art (LOEWE Start Professorship to M.N.H. and Excellence Program ‘The Adaptive Mind’) and the Deutsche Forschungsgemeinschaft under Germany’s Excellence Strategy (EXC 3066/1 ‘The Adaptive Mind’, 533717223). We thank Burkhard Maess, Yvonne Wolff-Rosier and Olaf Hauck for support and advice on all the MEG-related aspects of this project. We thank Ahmet Çeşmeci for assisting with computing the peak saccade curvature timing.

## Notes

### Competing Interest Statement

The authors have declared no competing interest.

### Summary of Updates

The whole manuscript has been revised substantially, with new analyses, an update of all figures and a major update of the text.

